# Deep learning of dynamic signatures resolves mechanistic ambiguity in complex signaling networks

**DOI:** 10.64898/2026.08.04.742863

**Authors:** Mustafa Ozen, Chaitra Agrahar, Francesca Zappa, Simone Bianco, Diego Acosta-Alvear, Carlos F. Lopez

## Abstract

Sparse experimental data often yields vast mechanistic hypothesis spaces with numerous equally probable models. Traditional model selection metrics like the Akaike Information Criterion fall short because they reduce models’ nonlinear dynamics to a scalar score that masks crucial mechanistic details. Here, we introduce an AI-driven framework treating model dynamics as learnable signatures. Using deep learning autoencoders, we embed the dynamic signatures of thousands of competing models into a low-dimensional latent space. Iterative clustering and physiological constraints systematically refine this space to a manageable number of testable hypotheses. Applying this to the Integrated Stress Response, a fundamental cellular homeostasis mechanism, we converged on 12 compelling mechanistic hypotheses from over 12,000 candidates. Notably, this framework confidently rejected structure-derived kinetic assumptions regarding higher-order PKR activation that traditional metrics could not resolve. Ultimately, this method provides a rigorous, full-state alternative to scalar metrics, accelerating the discovery of driving principles in complex biological systems.

## Introduction

The robust yet flexible behavior of biological systems is controlled by complex, non-linear networks of molecular interactions that govern processes from signaling cascades to gene regulation (Bonneau, 2008; Jordan et al., 2000). Gaining a mechanistic understanding of these complex systems is essential for identifying and precisely targeting the critical regulatory points within the network, which is fundamental for developing effective therapeutic strategies. Mechanistic models, often formulated as systems of ordinary differential equations (ODEs) or partial differential equations (PDEs), serve as powerful tools to represent hypothesized molecular interactions and predict system behavior (Albeck et al., 2008; Emadi et al., 2022; Jaqaman & Danuser, 2006; Ozen et al., 2020; Sevrin et al., 2024; Shockley et al., 2019). These models encapsulate the current understanding of a biological system, enabling quantitative predictions, facilitating the identification of critical regulatory nodes, and providing a framework for testing in silico hypotheses (Barh et al., 2014; Hat et al., 2016; Klein et al., 2022; Tuechler et al., 2025).

Despite their usefulness, the implementation and analysis of mechanistic models present significant challenges (Banga & Villaverde, 2025; Ozen et al., 2023; Tsopanoglou & del Val, 2021). The enormity of the biochemical space where molecular interactions occur results in an equally vast space for plausible mechanistic hypotheses. Every biological signaling pathway can involve several protein-protein interactions, post-translational modifications, and positive/negative feedback loops, each representing a potential model component (Aldridge et al., 2006). Traditionally, model construction relies on expert knowledge and iterative manual refinement, a process that can be time-consuming and prone to bias, potentially overlooking alternative valid explanations. Furthermore, the process of model calibration (fitting model parameters to experimental data) and subsequent model selection (choosing the most appropriate model that optimally balances predictive accuracy with biological validity) can be computationally intensive and statistically demanding, particularly when dealing with noisy, sparse, or high-dimensional biological data, which is common in -omics experiments typically used to build models (Alarid-Escudero et al., 2018; Irvin et al., 2023). Even for a single model structure with fixed hypotheses, ensuring parameter identifiability (the ability to uniquely estimate model parameters from data) is another critical obstacle that often limits confidence in mechanistic conclusions (Gutenkunst et al., 2007; Raue et al., 2009; Schaber and Klipp, 2011). Fitting a model with dozens of parameters to limited time-series data can lead to multiple parameter sets yielding similar fits, a common issue in complex biological systems where not all species or reactions are directly observable, making it difficult to discern the true underlying mechanism (Guillaume et al., 2019; Ortega et al., 2024). Therefore, in the face of these challenges, a robust and principled approach to model selection is essential to distinguish between competing hypotheses/parameter sets and ensure the biological relevance of the chosen model for downstream analyses.

Established approaches to model selection, such as the Akaike Information Criterion (AIC) and the Bayesian Information Criterion (BIC), provide statistical frameworks to balance model fit with complexity (Burnham & Anderson, 2002; Stoica & Selen, 2004). AIC estimates the out-of-sample prediction error, favoring models that generalize well, while BIC provides a larger penalty for model complexity, tending to select simpler models that are more likely to be accurate. While widely used, these approaches have limitations in the context of biological systems (Harbecke et al., 2024). They often rely on assumptions of normally distributed errors and large sample sizes, which are frequently violated in experimental biological datasets (Brewer et al., 2016). Their effectiveness can also be diminished when dealing with non-identifiable parameters or when the underlying data-generating process is non-linear, as is common in biological systems (Gu et al., 2018). Fundamentally, these criteria evaluate models strictly within the observable state space (i.e., experimentally measured species) and condense performance into a single scalar score. This inherently fails to capture the full spectrum of a model’s nonlinear dynamic phenotype, effectively rendering traditional metrics blind to the dynamics of unmeasured latent variables. Such a reductive approach can lead to inaccurate interpretations, as a model with a better score may not accurately reflect the underlying biological mechanism.

Artificial intelligence (AI) and machine learning (ML) have revolutionized diverse scientific disciplines by providing advanced capabilities for pattern recognition, predictive modeling, and automated decision-making. These computational methods are increasingly recognized for their potential to accelerate biological discovery and biomedical applications, ranging from target identification to precision drug design (Guglielmetti et al., 2025; Ocana et al., 2025; Visan and Negut, 2024; You et al., 2022). While AI/ML models are often perceived as “black boxes” due to their data-driven nature, their analytical power can be synergistically integrated with mechanistic modeling to overcome inherent challenges in hypothesis exploration. A prominent example of this synergy is Sparse Identification of Nonlinear Dynamics (SINDy), a framework that utilizes machine learning to extract sparse, explicit governing equations directly from time-series data (Brunton et al., 2016). By identifying the minimal set of nonlinear functions necessary to describe species derivatives, SINDy provides interpretable ODEs, effectively bridging the predictive accuracy of AI with the mechanistic clarity required for biological insight.

Here, we present a novel, AI-driven model selection workflow that moves beyond scalar metrics by treating model dynamics as a learnable signature. We leverage a deep learning autoencoder to embed the time-series trajectories of several competing mechanistic models, each with diverse structures and equivalent fits to experimental data, into a low-dimensional latent space. We then iteratively reduce the hypothesis space by clustering these dynamic embeddings and applying stringent biological constraints. We apply our approach to the Integrated Stress Response (ISR), a fundamental cellular homeostasis network, and we demonstrate how this component-by-component refinement converges on a manageable set of plausible mechanistic hypotheses from an initial pool of thousands.

## Results

### The Integrated Stress Response pathway

The Integrated Stress Response (ISR) is an evolutionarily conserved signaling network that allows cells to cope with stress and maintain homeostasis (Boone and Zappa, 2023; Costa-Mattioli and Walter, 2020). When stress is insurmountable, the ISR can switch from an adaptive repair state to a terminal programmed cell death state (Zappa et al., 2025). This dual role has implicated the ISR in aging (Kourtis and Tavernarakis, 2011) and various diseases, including cancer (Tian et al., 2021), neurodegeneration (Wolozin and Ivanov, 2019), and ischemic disease (Zhang et al., 2022).

The ISR (Figure 1) is governed by four stress sensor kinases (GCN2, HRI, PERK, and PKR) that detect amino acid shortages and ribosome collisions, low heme, disruption of proteostasis in the ER, and double-stranded RNA (dsRNA), respectively (Pakos-Zebrucka et al., 2016). Upon detection of their cognate stress inputs, each kinase homodimerizes and undergoes autophosphorylation, resulting in activation. The active kinases converge by phosphorylating the α subunit (eIF2α) of the heterotrimeric GTPase eIF2. Phosphorylated eIF2α inhibits its guanine nucleotide exchange factor, eIF2B (Clemens et al., 1982; Gordiyenko et al., 2019; Krishnamoorthy et al., 2001). In the absence of stress, eIF2, together with GTP and the initiator methionyl tRNA (Met-tRNAi), forms the eIF2-GTP-Met-tRNAi ternary complex (TC), which is required for translation initiation. The interplay between eIF2α phosphorylation and uninhibited eIF2B determines the levels of TC. Therefore, accumulation of phosphorylated eIF2α lowers TC availability and represses translation initiation, leading to a global repression of protein synthesis. Notably, these conditions simultaneously promote the selective translation of key transcription factors, such as ATF4 and CHOP, through a well-studied mechanism involving regulatory upstream open reading frames (uORFs) located in the 5’ region of their respective mRNAs (Harding et al., 2000; Hinnebusch et al., 2016). An ISR-embedded negative feedback loop is engaged by the induction of the phosphatase-regulatory subunit GADD34, which dephosphorylates eIF2α, restoring TC levels and terminating the signal (Connor et al., 2001; Novoa et al., 2001). Under severe stress, the ISR can switch to engage a terminal state triggered by the ISR transcription factor CHOP, and its downstream target, the death receptor DR5 (Zappa et al., 2025). Through these mechanisms, the ISR controls cellular adaptability or cell death (Costa-Mattioli and Walter, 2020; Holcik and Sonenberg, 2005; Hotamisligil and Davis, 2016).

**Figure 1:**
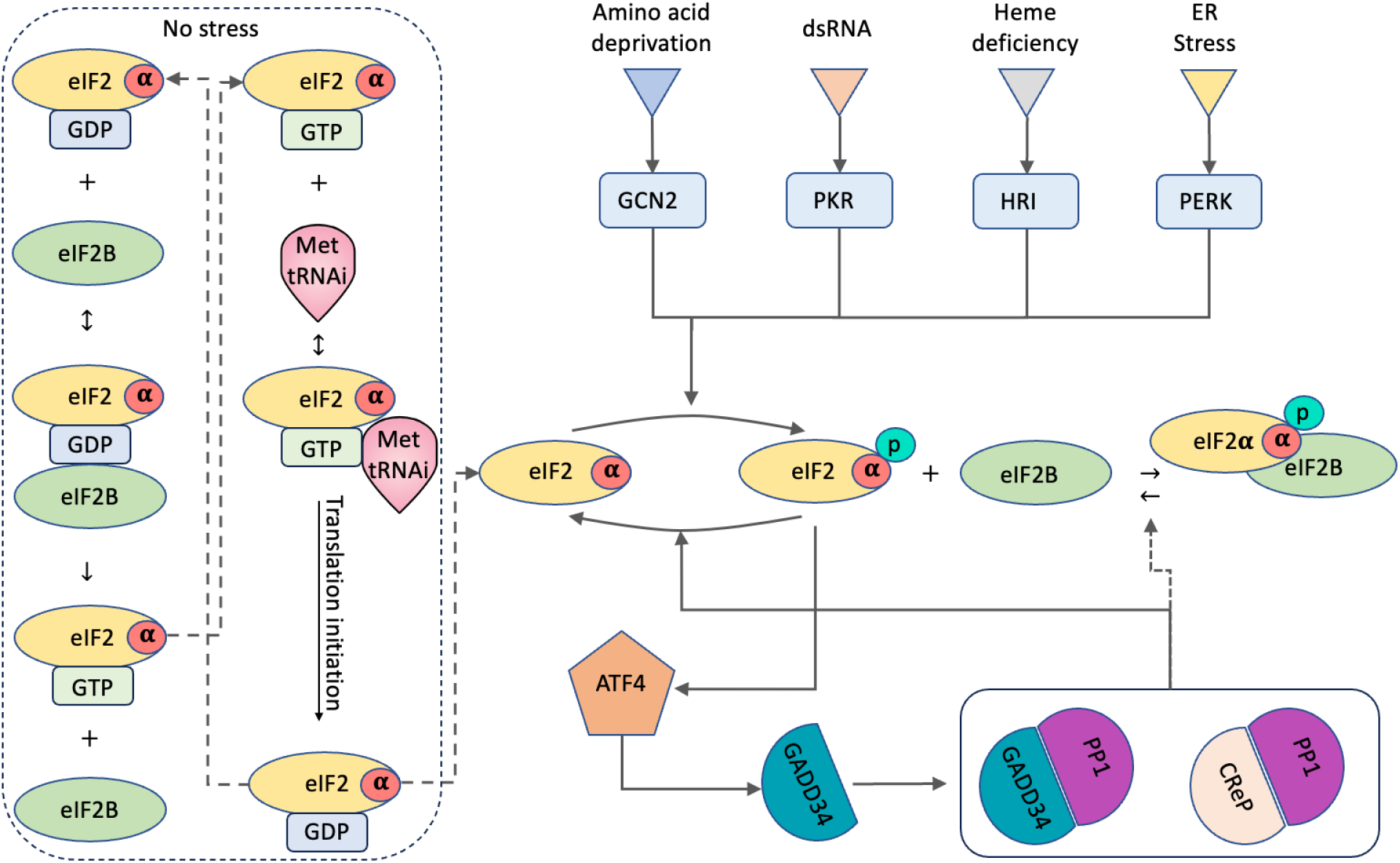
ISR in a nutshell. Model components at baseline and upon ISR activation. When no stress is detected, eIF2B hydrolyzes eIF2-GDP to eIF2-GTP, which then, together with Met t-RNAi, forms a ternary complex and initiates translation. Once any of the kinases senses a stressor, the active kinase phosphorylates the α subunit of eIF2 (eIF2α), leading to ATF4 expression, following induction of a negative feedback through GADD34 expression.

### Mechanistic modeling of the ISR and model calibration

Mechanistic models provide a quantitative framework to understand complex, nonlinear dynamics, such as those of the ISR. However, traditional modeling efforts often rely on a single, fixed structure, which may not capture the full range of plausible mechanisms supported by sparse experimental data. Existing mathematical models of the ISR (Batjargal et al., 2023; Klein et al., 2022; Pontisso et al., 2023) illustrate this challenge: they are often tailored to specific biological contexts or purposes, making it difficult to determine how well they represent the broader ISR molecular network across different cell types and stressors. Rather than seeking a single, universal model, a more rigorous approach involves evaluating a broad space of competing mechanistic hypotheses, ranging from varying kinetic laws (e.g., mass-action vs. Hill functions) to the presence of specific feedback loops. This results in a large ensemble of thousands of potentially valid explanations, requiring a principled methodology to identify the most biologically plausible hypotheses supported by the available data.

To systematically explore the vast mechanistic space of the ISR, we first divided the network into multiple functional modules (Figure 2A). For each module, we compiled a set of plausible competing mechanistic hypotheses (encompassing various kinetic laws, reaction stoichiometries, and potential feedback loops) based on expert knowledge and literature review (Section A, Supplementary Document). We then leveraged a Julia-based modeling framework, Catalyst (see Methods), to combinatorially generate an extensive set of models. Each model represented a unique combination of one hypothesis selected from every module. This systematic approach yielded a total of 12,096 distinct mechanistic models, formulated as systems of ODEs that vary significantly in structure, the number of species, and free parameters (Figure 2C-D). This ensemble includes “small models” that exclude the core translation initiation components (e.g., GDP-GTP hydrolysis by eIF2B and TC formation, Figure 1) and “large models” that include them (Figure 2B).

**Figure 2:**
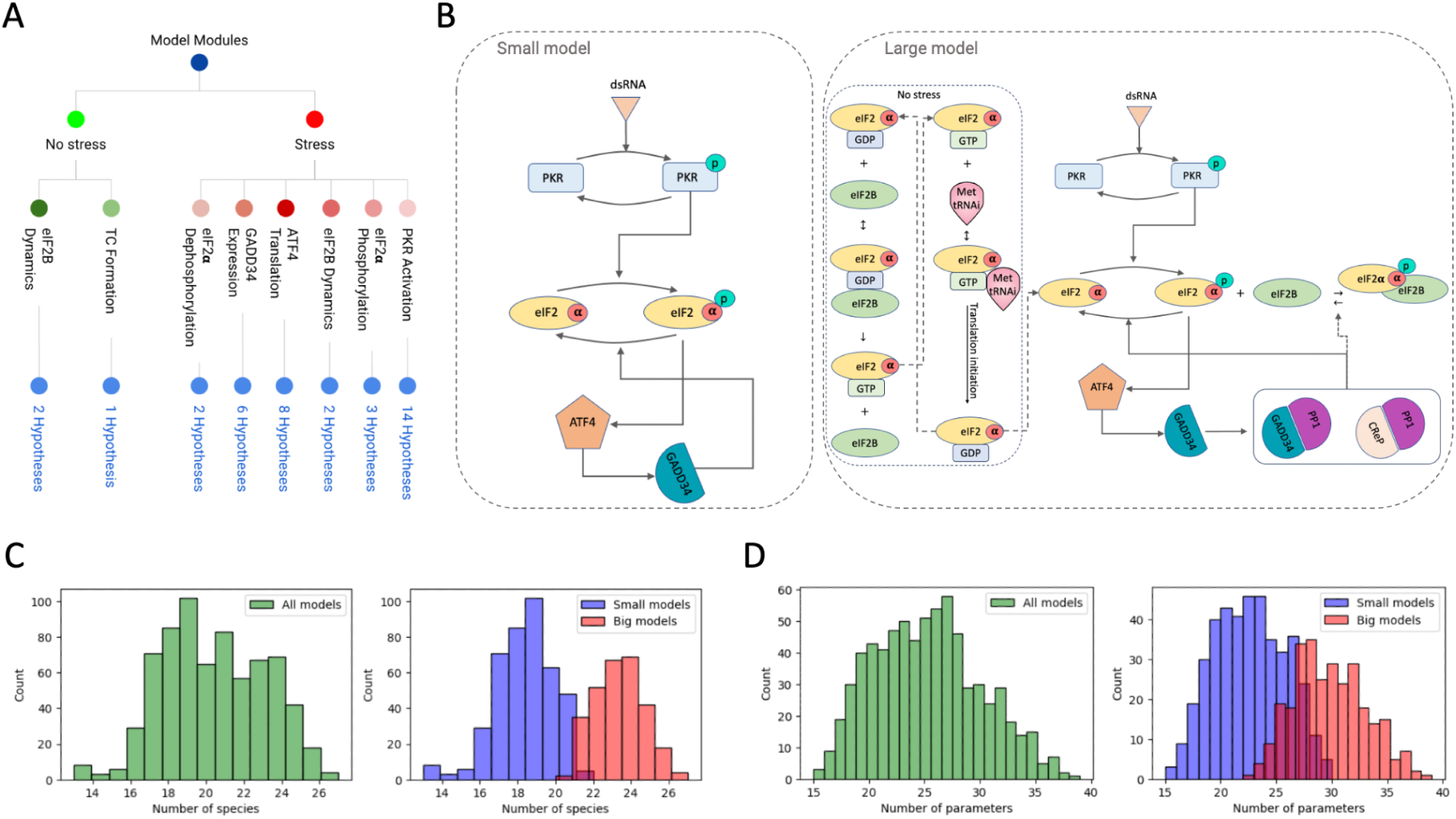
ISR hypothesis generation and modeling framework. (A) Modularized ISR processes. (B) Examples of generated models differ both in structure and kinetics. Some models do not include the “no stress” modules, which we refer to as “small model”, and some models include the “no stress” module, which we refer to as “large model”. (C-D) Distributions of the number of species and the number of parameters of the generated models.

To begin eliminating suboptimal models, we first calibrated the free parameters of each model using time-resolved proteomics data of Total PKR, ATF4, and GADD34 from quantitative proteomics measurements (Ozen et al., 2025). The data were obtained upon activation of a validated, synthetic ISR kinase, FKBP-PKR, which we have used in previous studies (Zappa et al., 2022; 2025). FKBP-PKR can be selectively activated using a drug-like small molecule, resulting in the induction of a “pure” ISR without potential confounder effects (Zappa et al., 2022). The data included 8 time points: 0, 1, 2, 4, 6, 8, 16, and 24 hours of FKBP-PKR activation in H4 neuroglioma cells (Ozen et al., 2025).

The aforementioned calibration process identified 709 out of 12,096 models that fit the proteomics data equally well and fall under a predetermined minimum-squared error (MSE) threshold, which we identified based on model divergence from experimental data (Figure 3A-B). This initial screening successfully eliminated certain hypotheses associated with specific ISR modules (see Supplementary Document), such as those related to the PKR module, while highlighting others that appeared more frequently across the good-fitting models (Figure 3C). Despite their comparable agreement with the measured data, these models exhibited significant divergence in the simulated dynamics of non-experimentally measured components, as evidenced by their high standard deviations (Figure 3D). Therefore, it is not possible to accurately select a single representative model through scalar-based selection criteria that solely consider model complexity and fit, and can be misleading (as elaborated below). Accurate model selection, therefore, requires factoring in the complex, nonlinear dynamics of the entire system.

**Figure 3:**
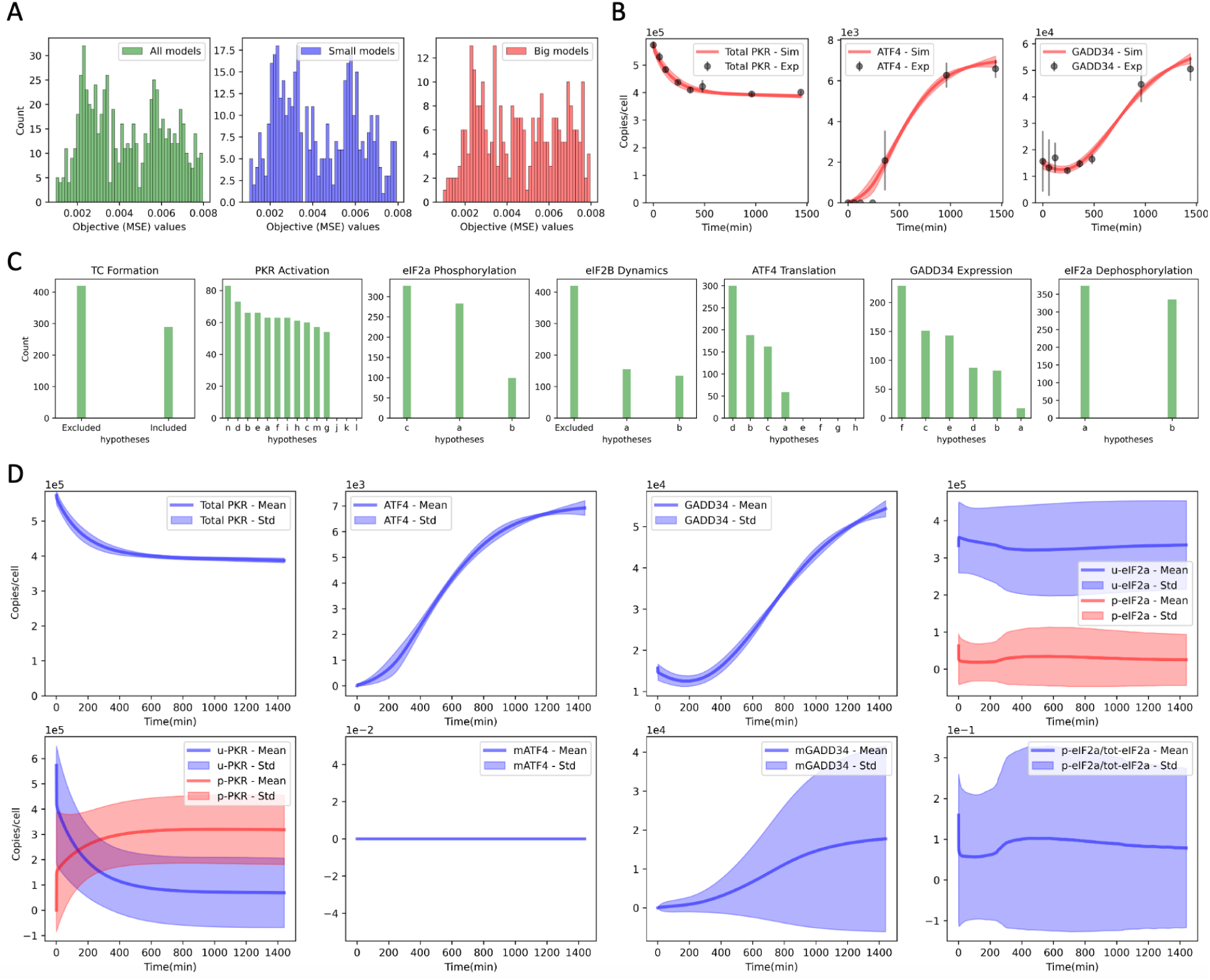
ISR hypothesis generation and modeling framework. A) MSE distributions of 709 good-fitting models. MSE distributions of all 12,096 models are provided in Figure S8. (B) 709 models’ goodness of fit (MSE < 0.008, see Methods). (C) Hypotheses distributions in good-fitting models. (D) 709 good-fitting models’ overall dynamics, including the components that were not experimentally measured. Notice that although all 709 models have equally good fits to PKR, ATF4, and GADD34 measurements, they exhibit distinct dynamics for the other components, such as u-PKR and p-PKR. Therefore, further model selection is needed for downstream conclusions.

### Traditional scalar metrics fail to capture dynamic signatures

To establish a baseline for model selection, we first applied traditional, scalar-based information criteria to our ensemble of 709 calibrated models. Specifically, we evaluated the Akaike Information Criterion (AIC) and its corrected version (AICc) for each model to identify candidates that optimally balance experimental goodness-of-fit with structural parsimony. As expected, AIC and AICc heavily penalized highly parameterized networks, exclusively isolating three structurally simple models as the top candidates. Upon structural inspection, all three of these AICc-selected models completely omitted the dynamics of TC and eIF2B (Figure 4), rewarding parsimony above all else.

**Figure 4:**
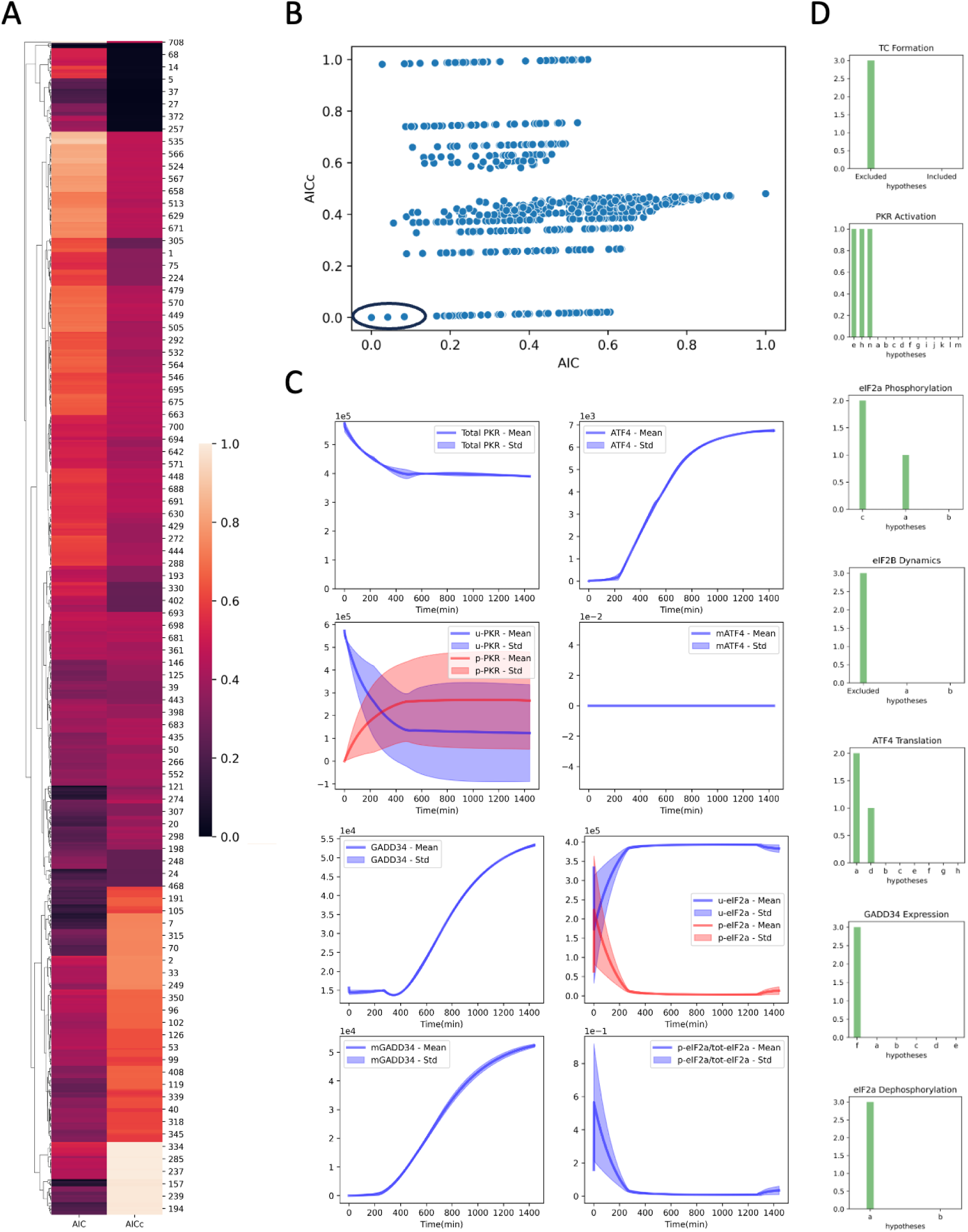
Model selection via traditional scalar-based scoring (AIC and AICc). Traditional scalar metrics fail to capture full-state dynamic topology. (A) AIC and AICc scores evaluate models strictly on observed data. (B) The three models selected by AIC/AICc reward parsimony but ignore unmeasured state spaces. The intersection of lowest AIC and AICc scores (circled) was evaluated to filter out mathematical artifacts associated with small-sample penalties (i.e., cases when number of parameters > number of samples, resulting negative penalty in AICc). (C) Consequently, AIC-selected models generate highly non-physiological trajectories for latent variables, demonstrating the vulnerability of scalar metrics to partial observability. D) Hypotheses in the selected 3 models.

However, when we simulated the full-state trajectories of these top-scoring models, they produced biologically unreasonable predictions for latent, non-measured components. They exhibited highly non-physiological trajectories, such as the rapid, unconstrained phosphorylation and dephosphorylation cycling of eIF2α despite the presence of a sustained stress input (Figure 4C). This stark disconnect highlights a fundamental limitation of traditional scalar metrics: by reducing a complex, nonlinear dynamic phenotype to a single score evaluated solely on a few observable states, AIC and AICc act completely blind to the dynamic realism of the unmeasured system. Consequently, relying solely on these traditional criteria would have resulted in the selection of biologically flawed mechanisms while erroneously discarding a wide range of viable, dynamically robust hypotheses. This failure demonstrates that scalar metrics are insufficient for capturing full-state dynamic behaviours, necessitating a selection framework capable of evaluating the holistic dynamic signature of the entire network.

### Model selection via autoencoders

To further refine model selection, we subjected the 709 models that passed our initial filters to triage using autoencoders, a type of deep neural network used in unsupervised machine learning that encodes input data into a lower-dimensional representation and then decodes it to reconstruct the original input, to enable dimensionality reduction, feature extraction, and anomaly detection in linear and nonlinear data (Bank et al., 2023). This approach enabled the unearthing of latent, lower-dimensional representations of each model’s dynamic signature.

We focused on a highly ambiguous component of the models (i.e., a species whose trajectories have high StDev in equiprobable models), specifically the concatenated unphosphorylated-PKR (u-PKR) and phosphorylated-PKR (p-PKR) trajectories (Figure 5A) to train an autoencoder (see Methods). We clustered the 709 models in the latent space based on their embedded dynamic signatures, resulting in four clusters representing distinct dynamic behaviors of PKR activation (Figure 5B). To identify the most biologically plausible behavior from the latent manifold, we applied two strict physiological prior constraints derived from established literature with supporting experimental data: (1) a gradual activation profile requiring a monotonic increase in p-PKR over the initial phase (Klein et al., 2022; Zappa et al., 2022), and (2) a progressive accumulation of p-PKR as a function of input level (Ozen et al., 2025). This approach narrowed the model space to 364 models (green cluster in Figure 5B) that satisfied both criteria. Other model clusters were discarded as they represented behaviors that did not align with our two prior-knowledge-based criteria; for instance, the rapid saturation of p-PKR (orange cluster) or a low activation profile (red cluster) deviated significantly from the expected gradual and dose-dependent accumulation of p-PKR.

**Figure 5:**
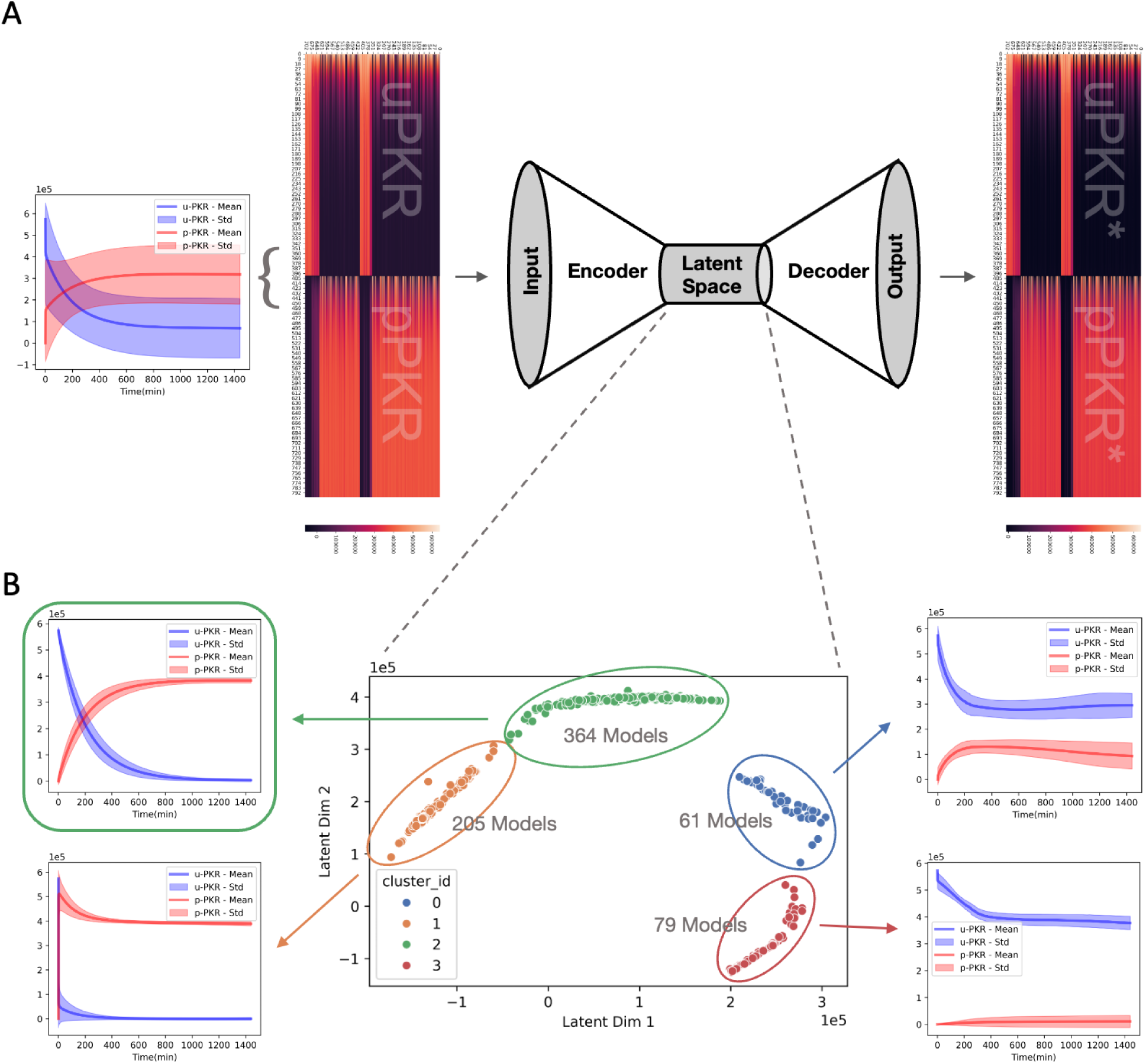
Model selection via autoencoders: overview of a single iteration. (A) Embedding u-PKR and p-PKR dynamics to learn a latent representation. Given highly variant u-PKR and p-PKR dynamics, we concatenate them into a single vector per model (each column of the heatmap). Then, we train an autoencoder to learn a structured latent representation of the dynamics data with a goal of capturing distinct behaviors. (B) Latent space representations of the models based on concatenated u-PKR, p-PKR dynamics. Clustering the latent representations of the models identifies unique u-PKR and p-PKR dynamics. With the expert and prior knowledge, we select a cluster (the green cluster in this case) and move on to the next iteration.

The selected 364 models exhibited significantly different dynamic behaviors in other non-measured components, including unphosphorylated-eIF2α (u-eIF2α) and phosphorylated-eIF2α (p-eIF2α), major ISR output determinants that lie immediately downstream of active, p-PKR. To find similarities at this level of the signaling cascade, we next trained an autoencoder using the concatenated time-series trajectories of u-eIF2α and p-eIF2α obtained from each of the 364 models (Figure S1). Because the initial phosphorylation level of eIF2α ‘overshoots’ before stabilizing back into a new, lower steady-state level (Rajesh et al., 2015; Wek et al., 2023) due to GADD34 induction (Connor et al., 2001; Costa-Mattioli and Walter, 2020), we selected the models based on this dynamic criterion in the next iteration of the model selection. This selection yielded 40 models (red cluster in Figure S1B) whose dynamics correctly captured the p-eIF2a ‘overshoot and adaptation’ signature. Moreover, the models in this cluster exhibit u-eIF2α and p-eIF2α activity profiles that correlate with u-PKR and p-PKR activities. Lastly, we refined this process by training a new autoencoder for the p-eIF2α to total eIF2α ratio over time (Figure S2), which eliminated an additional 28 models by applying a strict, empirically validated physiological threshold. Concurrent biochemical validation confirms that ATF4-dependent ISR signaling operates as a switch strictly licensed when the p-eIF2α to total eIF2α ratio reaches approximately 30% (Klein et al., 2022; Ozen et al., 2025). Models falling outside the latent clusters that satisfy this empirically validated 0.3 threshold were systematically discarded. These sequential iterations ultimately resulted in a final set of 12 models (Figures 6, 7, S3, Table S1) that holistically represent biologically reasonable dynamics. We chose not to cluster the models further based on GADD34 mRNA trajectories because the observed high StDev is an artifact of the model design itself. This variance arises because the initial hypotheses for the GADD34 expression module were deliberately structured to test different levels of model complexity, meaning some models explicitly included GADD34 mRNA as a species while others did not, making its trajectory unsuitable for final dynamic clustering.

**Figure 6:**
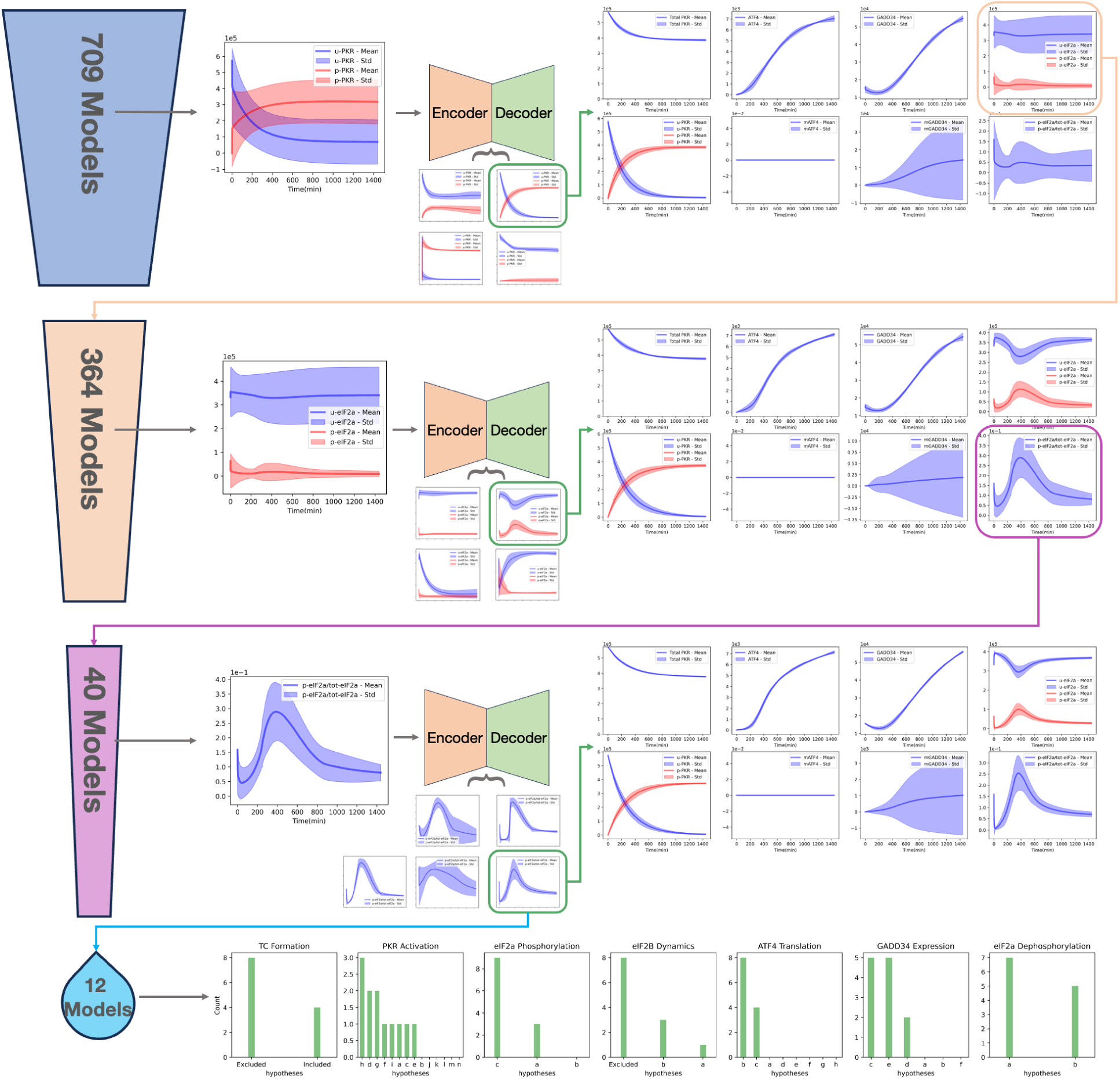
Iterations of model selection via autoencoders. Starting with 709 models, iteratively training an autoencoder, clustering the latent space, and selecting a subcluster per model component based on established physiological constraints yields 12 models with the most relevant overall dynamics.

### Characterization of selected models

A more detailed analysis of the individual modules within the 12 aforementioned models revealed that the inclusion or exclusion of the TC and eIF2B dynamics was represented with similar frequency. This finding demonstrates that complex models (i.e., those that explicitly include TC and eIF2B dynamics) and simpler models (i.e., those that exclude them) can equally capture ISR behaviors. As established earlier (Figure 4), traditional scalar metrics like AIC and AICc improperly rejected this entire class of complex, viable models in favor of structurally sparse networks that failed to capture physiological latent dynamics. By contrast, our deep learning framework successfully retained these dynamically valid complex models alongside their simpler counterparts, proving that full-state dynamic embedding preserves the dynamic realism that is required for meaningful mechanistic insight.

In the PKR module, all hypotheses incorporating more complex PKR activation mechanisms, such as explicit dimerization and self-activation kinetics (hypotheses k to n in Section A, Supplementary Document), were eliminated by our autoencoder-based approach. This result suggests that a simpler, reversible mass-action formalism of the reaction governing the phosphorylation of u-PKR to its active form, p-PKR, is sufficient to recapitulate the observed system dynamics. Nevertheless, the selected models remain inconclusive regarding the precise dynamics of PKR activation by stress (i.e., the forward reaction rate) and the potential negative feedback from downstream signaling that could reverse this reaction rate (i.e., the final models exhibit significant variation in these parameters). To distinguish among the remaining mechanistic hypotheses for the PKR module, additional experimental data, particularly high-resolution phospho-proteomics measurements, are required. Importantly, our screening process allowed us to test and eliminate a specific mechanistic hypothesis—the structurally derived assumption (Dar et al., 2005) that a single active PKR dimer phosphorylates two eIF2 molecules simultaneously (hypothesis b), which is discussed in detail in the next section.

In addition, most of the models in our final set rely on a basal eIF2α phosphorylation mechanism (hypothesis c in the eIF2α phosphorylation module) to accurately recapitulate the dynamics of p-eIF2α, a finding that is consistent with established literature (Lu et al., 2004). The inclusion of this basal activity is essential for model function, as it provides a necessary non-zero starting point for the active signaling cascade. This baseline activity ensures the system is poised for rapid response to acute stress inputs, properly licensing the subsequent accumulation of p-eIF2α and the downstream translation of ATF4, thereby preventing the kinetic lag times beyond physiological expectations (discussed in Ozen et al., 2025) that would otherwise violate the response observed experimentally. This outcome reinforces the utility of our AI-driven framework in selecting models that not only fit the endpoint data but also adhere to the subtle yet fundamental mechanistic requirements of the ISR system.

As observed for PKR, the selected models also favored simpler mechanistic hypotheses for the ATF4 and GADD34 modules, unencumbered by ATF4 mRNA dynamics. Only two out of eight hypotheses for ATF4 translation occurred in the selected models, and both of them consider a forward reaction rate dependent on either the p-eIF2α to total eIF2α ratio, or the p-eIF2α to u-eIF2α ratio, both of which represent inherently similar dynamics. This result strongly suggests that the p-eIF2α to total eIF2α ratio controls the translation of ATF4, corroborating decades of experimental data on ATF4 induction during the ISR. This result indicates that the cell does not simply measure the raw absolute amount of p-eIF2α to induce the ISR, but instead relies on a ratiometric measurement correlated with the sensing of stress and ISR induction. Indeed, in recent work, we predicted and experimentally validated that the p-eIF2α to total eIF2α ratio licenses ISR activation (Ozen et al., 2025). Beyond successfully recovering these validated biological constraints, our dynamic embedding approach also demonstrated the capacity to actively interrogate and filter out conflicting mechanistic assumptions.

### Dynamic embedding refutes structurally derived PKR higher-order activation kinetics

A major advantage of evaluating the full dynamic topology is the ability to resolve specific mechanistic controversies. Based on established structural data (Dar et al., 2005), it has been hypothesized that a single active PKR dimer can phosphorylate two eIF2 molecules simultaneously (hypothesis b). However, all hypotheses incorporating these more complex, higher-order PKR activation mechanisms were eliminated by our autoencoder-based approach. This demonstrates that a simpler, reversible reaction governing the phosphorylation of u-PKR to p-PKR is mathematically sufficient, and that enforcing higher-order substrate recognition breaks the temporal topology of the full system. Crucially, this AI-driven rejection of cooperative PKR kinetics is empirically supported by independent biochemical assays. Dose-response immunoblots of FKBP-PKR activation demonstrate that p-eIF2α accumulates progressively alongside active p-PKR, lacking the steep, ultrasensitive transition that would mathematically obligate the 1:2 enzyme-substrate stoichiometry postulated by structural models (Ozen et al., 2025). Thus, the latent space embedding correctly vetoed a local structural assumption by enforcing global kinetic consistency. As visualized in the final mechanistic wiring diagram (Figure 7), all models enforcing complex self-activation kinetics (PKR hypotheses k through n) and higher-order substrate recognition (eIF2Phos hypothesis b) were entirely pruned from the latent manifold.

**Figure 7:**
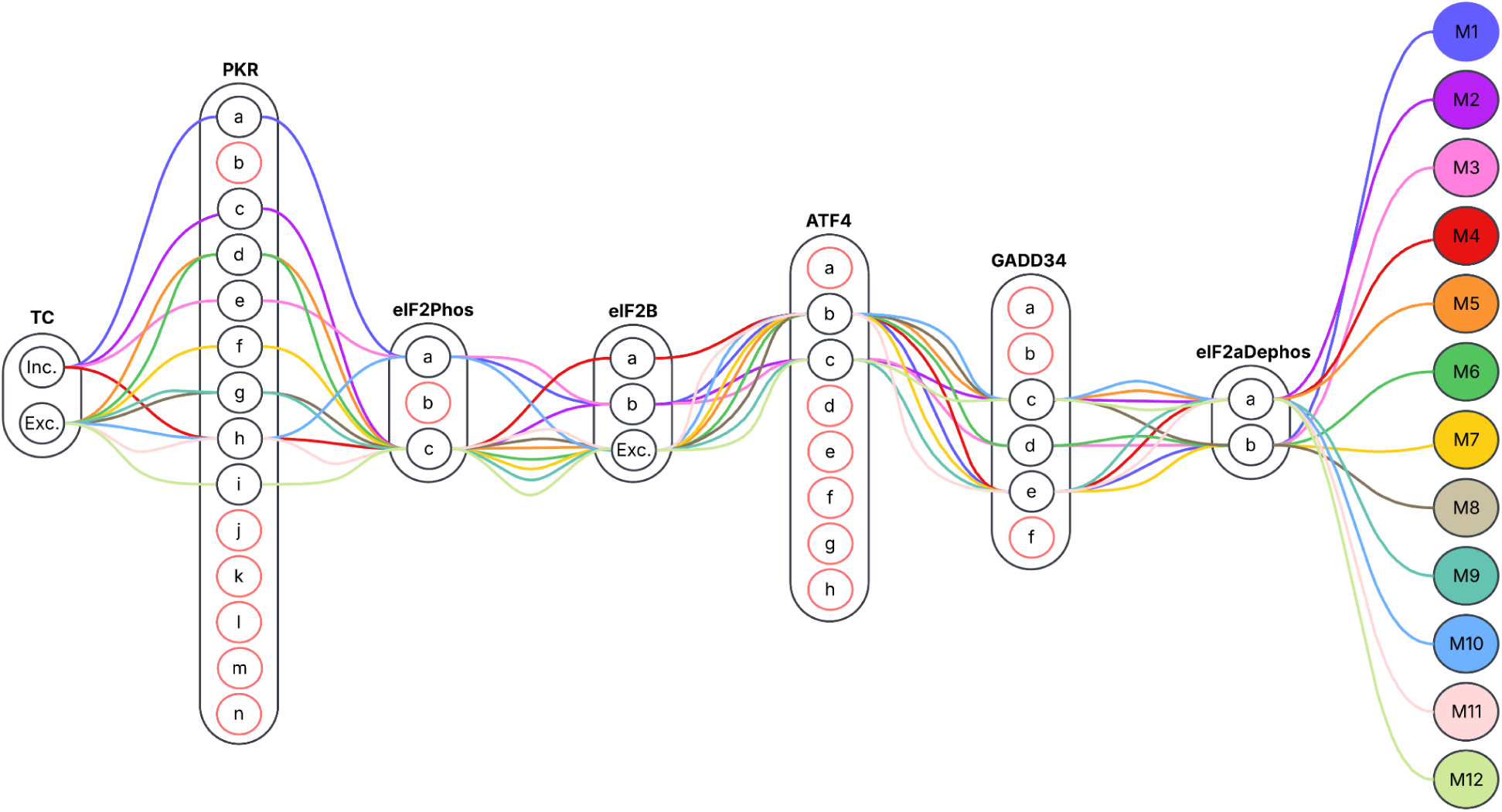
Wiring of the selected 12 models in the hypotheses space. Each letter represents a hypothesis for the associated ISR module (listed in the Supplementary Document). All the red-circled hypotheses were eliminated either during model calibration or autoencoder-based model selection

### Robustness of model selection

To further validate our approach, we conducted a parallel analysis in which we varied the sequence of components that yielded the aforementioned 12 models. Our primary analysis proceeded via the sequential embedding of the dynamics of u-PKR and p-PKR, followed by u-eIF2α and p-eIF2α, and finally the p-eIF2α to total eIF2α ratio. We then tested the robustness of this procedure by using an alternative sequence, starting with the p-eIF2α to total eIF2α ratio, then u-PKR and p-PKR, and concluding with u-eIF2α and p-eIF2α (Figures S4-S6). This alternative ordering resulted in a final set of 17 models, which included all 12 models obtained in our initial analysis (Figure S7). This demonstrates that our deep learning framework identifies a stable, underlying ‘mechanistic manifold’ that represents the true topological structure of the system, rather than a path-dependent artifact of the clustering sequence. By confirming that the identified set of hypotheses is robust and independent of the arbitrary sequence of components used for iterative refinement, we validate the mathematical objectivity of the selection pipeline.

## Discussion

The systematic exploration of mechanistic hypotheses for complex biological systems remains a challenge in computational biology. The combinatorial nature of molecular interactions and kinetic laws invariably generates a vast hypothesis space, where numerous models can equally reproduce observed experimental data through fundamentally different underlying mechanisms. Our work addresses this challenge by introducing a novel, AI-driven framework for mechanistic model selection that bypasses the limitations of traditional, scalar-based approaches. By leveraging deep learning autoencoders to embed and cluster models based on their dynamic signatures, we provide a principled method for navigating and reducing an immense hypothesis space.

The efficacy of our approach is demonstrated through its application to the ISR. In this setting, we successfully reduced a starting pool of equiprobable 12,096 combinatorially generated models (Figure 2) to a final set of 12 (Figure 7) that are consistent with experimental data and biological knowledge. A key finding from our analyses is that simpler models, which exclude TC dynamics, and more complex models, which include them, were found to be equally plausible. This highlights a critical limitation of traditional model selection criteria, such as AIC and BIC. Because these metrics rigidly penalize parameter complexity, they systemically discarded the entire subset of valid, complex mechanistic explanations in our ensemble. Our findings emphasize that while scalar metrics are useful for enforcing strict parsimony, they are inadequate for biological systems where complex topological features, such as TC and eIF2B dynamics in ISR, must be evaluated holistically to maintain full-state dynamic signatures.

The necessity of full-state dynamic embedding is further underscored by extreme non-linear behaviors inherent to the ISR. Recent empirical work has identified a biphasic ‘translational cliff’, a regime where saturating p-eIF2α fully depletes ternary complexes, causing a paradoxical collapse in ATF4 translation at high stress amplitudes (Ozen et al., 2025). Because our models were calibrated to single-dose data, traditional scalar metrics like AIC only evaluate fitness within that restricted observable regime, effectively remaining blind to out-of-sample multi-dose phenomena like the translational cliff. AIC cannot logically penalize models for failing to predict a boundary absent from the training data. By contrast, embedding the full dynamic signature into a latent manifold groups models by their inherent topological behaviors across the entire unmeasured state space. This structural grouping allows us to actively query the latent space and isolate models whose underlying wiring is capable of accommodating these extreme physiological boundaries, even when trained on sparse data.

Our iterative selection process provided specific mechanistic insights into the ISR. Specifically, our models determined that a less complex, reversible reaction is sufficient to explain PKR activation, eliminating hypotheses that included explicit dimerization and self-activation kinetics. Our models also strongly support the inclusion of the basal eIF2α phosphorylation mechanism, a finding that aligns with the literature (Lu et al., 2004). Moreover, the models also provide strong evidence that the p-eIF2α to total eIF2α ratio is a key determinant of ATF4 translation, a hypothesis that we recently validated experimentally (Ozen et al., 2025). Notably, the framework also allowed us to test and reject the hypothesis that a single active PKR can phosphorylate two eIF2 molecules, one per promoter, a kinetic assumption reasoned from the established dimeric structure and higher-order substrate recognition reported previously (Dar et al., 2005). These findings exemplify the method’s ability to not only filter out hypotheses but also to scrutinize and refine specific kinetic assumptions carefully.

Beyond the specific insights into the ISR, our framework represents a new paradigm to navigate hypothesis uncertainty in any complex biological system. This expert-guided AI approach is readily generalizable to other fields where a multitude of mechanistic hypotheses can explain sparse experimental data. For example, it could be used to resolve competing models of drug resistance in cancer signaling pathways or to identify the most plausible network topologies in immunology or metabolic flux. By treating model behavior as a learnable dynamic phenotype, our method transforms model selection from a simple scoring exercise into a data-driven process that reveals the fundamental operating principles of complex biological systems, promising to accelerate biological discovery.

### Limitations

Our methodology has inherent limitations. First, selecting the most reasonable cluster in the latent space relies on an “expert in the loop” to interpret the biological relevance of the embedded dynamics, which prevents the full automation of the selection process. Second, the method’s effectiveness is contingent on the quality and breadth of the initial experimental data used for model calibration; poor and limited data can lead to high uncertainty in the identified good-fitting models and their parameters. Third, the scanning of the parameter space is limited by the computational tractability of model calibration and the sheer size of the hypothesis/model space. Fourth, structural and kinetic differences among the candidate models generate diverse systems of ODEs, some of which are challenging for standard ODE solvers. Fifth, even though we utilized a metaheuristic optimization approach, namely Simulated Annealing, to identify the parameters of good-fitting models, high-dimensional parameter spaces are notoriously plagued by local minima. It is possible that some models were discarded not because their structural hypotheses were invalid, but because the algorithm failed to find the global optimum within the allotted 100,000 iterations. Thus, our initial pool of 709 good-fitting models represents a robust sample of viable parameterizations rather than an exhaustive catalog of all possible solutions. An ideal approach would involve a Bayesian parameter inference scheme to identify the posterior distributions of the parameters. However, running such an inference for even a single complex model remains a significant challenge for high-dimensional nonlinear problems (Linden et al., 2022). Sixth, the iterative, component-by-component application of autoencoders to refine the hypothesis space is time-consuming. Instead, training a single autoencoder on the concatenated dynamics of all model components simultaneously could allow a holistic assessment of each model’s behavior. However, such an approach would not be free from significant challenges, including the need for a much larger neural network with a substantially greater number of parameters, which demands more computing power and time. This limitation notwithstanding, our iterative approach is more effective at discerning subtle yet mechanistically important differences in models’ dynamic signatures (e.g., small variations in amplitude or transient signaling kinetics) that a single, all-encompassing autoencoder might overlook. Finally, we utilized K-Means clustering for latent space discretization. While effective as a heuristic for this initial framework, deep autoencoder embeddings often form complex, non-Euclidean manifolds. Future iterations of this pipeline will benefit from implementing density-based topological clustering algorithms, e.g., DBSCAN (Ester et al., 1996), to map these latent spaces without assuming spherical cluster geometries.

## Methods

### Mechanistic model implementation

To systematically explore the vast hypothesis space of the ISR pathway, we first modularized the network into seven key components: TC formation, PKR activation, eIF2α phosphorylation, eIF2B dynamics, ATF4 translation, GADD34 expression, and eIF2α dephosphorylation. For each of these modules, we curated a set of competing hypotheses, varying in both complexity and kinetics (Section A, Supplementary Document). Considering all possible combinations of these hypotheses across the modules, we combinatorially generated a total of 12,096 distinct mechanistic models (e.g., a system of ODE represented by the sequence TC formation = “included”, PKR activation = “b”, eIF2α phosphorylation = “c”, eIF2B dynamics = “a”, ATF4 translation = “b”, GADD34 expression = “c”, eIF2α dephosphorylation = “b” forms a model).

The models were implemented in the Julia programming environment using the Catalyst modeling package (Loman et al., 2023). Julia was selected for its superior computational efficiency, particularly for biological applications (Pal et al., 2024), and for its access to a suite of stable and performant ODE solvers. We utilized the AutoVern9 with Rodas5p solver, which automatically adjusts for system stiffness, to ensure accurate and robust numerical solutions for the diverse range of model kinetics. This meticulous implementation strategy was an essential prerequisite for the subsequent large-scale model calibration and selection process.

### Mechanistic model calibration

Initial conditions for each model’s species were set based on corresponding experimental measurements provided in Ozen et al. (2025). The calibration of the models’ free parameters was performed using a metaheuristic optimization approach, specifically the Simulated Annealing algorithm from Julia’s optimization package, with a maximum of 100,000 iterations to ensure a robust search of the parameter space. We arbitrarily initialized model parameters, considering larger/smaller values for binding/unbinding reactions. The parameter scan was done on a log scale, and the range [-4, 4] was searched for all parameters. The objective was to minimize the total mean squared error (MSE) between the simulated and experimental time-series data for Total PKR, ATF4, and GADD34, as defined by the following objective function:

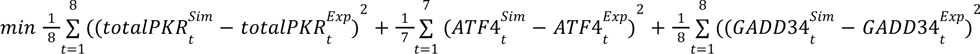

Note that the ATF4 measurement includes seven time points due to the exclusion of measurement at the 8^th^ hour as discussed in Ozen et al. (2025). To enhance the efficiency of the search space scanning and accelerate convergence, all experimental data and corresponding simulation outputs were min-max normalized, thereby eliminating the effects of different scales of protein abundances. This particular calibration strategy allowed us to identify the initial pool of 709 models that demonstrated an equally good fit to the experimental data.

709 out of 12,096 models that fit the proteomics data equally well were selected by thresholding models based on their MSE values. MSE = 0.008 was selected based on models’ fitness divergence from data as MSE increases. MSE > 0.008 results in models with worse fitness to at least one of the objective molecules (PKR, ATF4, GADD34).

### Autoencoder implementation and training

The autoencoder was implemented in Python using the PyTorch package. The encoder and decoder were structured as follows: the input layer (400 nodes) → hidden layer 1 (200 nodes) → hidden layer 2 (100 nodes) → latent space (2 nodes) → hidden layer 3 (100 nodes) → hidden layer 4 (200 nodes) → output layer (400 nodes). We fixed the activation functions of all layers to ReLU (rectified linear unit), except for the last layer, which was set to either ReLU or Sigmoid, depending on the species’ dynamics range. The dynamics of each input species were uniformly sampled at 400 time points. Note that the time course sampling rate is a hyperparameter that we did not thoroughly experiment with. Having more samples will surely be beneficial, as it increases the likelihood of capturing more information about nonlinear and perhaps transient dynamics. Conversely, having a lower number of samples may affect the structure of the latent space. The choice of 400 time points provided sufficient resolution to capture both fast transient kinetics and long-term steady-state behavior without introducing unnecessary computational overhead. The training of the autoencoder was run for 2000 epochs with a learning rate = 1e-4, an MSE loss function, and the Adam optimizer. Overall, we trained an autoencoder and learned a latent space representation in 2D per single model component. Visual inspection of the decoded trajectories and the low final MSE reconstruction loss confirmed that the 2D latent space successfully captured the fundamental temporal variance of the system without significant information loss (Figures 5A, S1A, and S2A).

### Latent space clustering

To cluster the latent space model representations, we used K-Means clustering in the Python Scikit-Learn package (Pedregosa et al., 2011). We computed distortion versus different K values to identify the optimal number of clusters per iteration.

## Acknowledgments

We would like to thank Francesca Ratti for insightful conversations and critical feedback on this work.

## Data and Code Availability

The quantitative proteomics dataset used for model calibration is detailed in Ozen et al. (2025) and is available at the GitHub repository: https://github.com/altoslabs/isrmm. The custom Julia and Python scripts used for mechanistic model generation and latent space embeddings are available from the corresponding author upon reasonable request. Detailed mathematical formulations, module hypotheses, and neural network hyperparameters are provided in the Methods section and Supplementary Document to enable methodological replication.

