## Supplementary Document for "Deep learning of dynamic signatures resolves mechanistic ambiguity in complex signaling networks"

### A. List of hypotheses for each ISR module:

In the listed reactions below, ":" indicates complexation, "⇌" indicates reversible reactions, and "⟶" indicates forward reactions.

1. **Ternary Complex Formation:** If included in the models, we assume that the ternary complex is formed in one way.

   Met-tRNAi + u-eIF2a$_{GTP}$ ⇆ Met-tRNAi:u-eIF2a$_{GTP}$
   Met-tRNAi:u-eIF2a$_{GTP}$ → u-eIF2a$_{GDP}$ + Protein ($k_f = c_f$, $k_r = c_r$, $k_c = c_c$)

2. **PKR Activation:**
   a. $PKR_u \leftrightarrows PKR_p \ (k_f = 1,\ k_r = 0.1)$
   b. $PKR_u \leftrightarrows PKR_p \ (k_f = 1,\ k_r = \frac{ueIF2a}{tot-eIF2a}k)$
   c. $PKR_u \leftrightarrows PKR_p \ (k_f = 1,\ k_r = \frac{ueIF2a}{peIF2a}k)$
   d. $PKR_u \leftrightarrows PKR_p \ (k_f = 1,\ k_r = \frac{ATF4}{K + ATF4}k)$
   e. $PKR_u \leftrightarrows PKR_p \ (k_f = \frac{S}{K+S}k,\ k_r = 0.1)$
   f. $PKR_u \leftrightarrows PKR_p \ (k_f = \frac{S}{K+S}k,\ k_r = \frac{ueIF2a}{tot-eIF2a}k)$
   g. $PKR_u \leftrightarrows PKR_p \ (k_f = \frac{S}{K+S}k,\ k_r = \frac{ueIF2a}{peIF2a}k)$
   h. $PKR_u \leftrightarrows PKR_p \ (k_f = \frac{S}{K_1+S}k,\ k_r = \frac{ATF4}{K_2 + ATF4}k)$
   i. $PKR_u \leftrightarrows PKR_p \ (k_f = 1,\ k_r = \frac{PKRu}{PKRtot}k)$
   j. $PKR_u \leftrightarrows PKR_p \ (k_f = 1,\ k_r = \frac{PKRu}{PKRp}k)$
   k. $PKR_u + PKR_u + S \rightarrow PKR^* + S \ (k_f = \frac{PKRp}{PKRu}k)$
      $PKR^* + PKR_u + PKR_u \rightarrow 2PKR^* (k_f = \frac{PKRp}{PKRu}k)$
      $PKR^* \rightarrow PKR_u + PKR_u (k_r = \frac{ATF4}{K + ATF4}k)$
   l. $PKR_u + PKR_u + S \rightarrow PKR^* + S \ (k_f = \frac{PKRtot}{PKRu}k)$
      $PKR^* + PKR_u + PKR_u \rightarrow 2PKR^* (k_f = \frac{PKRtot}{PKRu}k)$
      $PKR^* \rightarrow PKR_u + PKR_u (k_r = \frac{ATF4}{K + ATF4}k)$
   m. $PKR_u + PKR_u + S \rightarrow PKR^* + S \ (k_f = (\frac{S}{K+S}k)\ )$

$PKR^{*} + PKR_{u} + PKR_{u} \rightarrow 2PKR^{*} (k_{f} = \frac{S}{K+S} k)$

$PKR^{*} \rightarrow PKR_{u} + PKR_{u} (k_{r} = \frac{ATF4}{K + ATF4} k)$

n. $PKR_{u} + PKR_{u} + S \leftrightarrows PKR_{u} : PKR_{u} + S \; (k_{f} = \frac{S}{K+S} k, \; k_{r} = 0.01)$

$PKR_{u} : PKR_{u} \rightarrow PKR^{*} (k_{f} = \frac{S}{K+S} k)$

$PKR_{u} + PKR_{u} + PKR^{*} \rightarrow PKR_{u} : PKR_{u} + PKR^{*} \; (k_{f} = \frac{S}{K+S} k)$

$PKR^{*} \rightarrow PKR_{u} + PKR_{u} (k_{r} = \frac{ATF4}{K + ATF4} k)$

3. **eIF2a Phosphorylation:**
   a. $PKR^{*} + ueIF2a_{GDP} \leftrightarrows PKR^{*} : ueIF2a_{GDP} \rightarrow PKR^{*} + peIF2a_{GDP}$
   $(k_{f} = 1, \; kr = 0.1, \; k_{c} = 1)$
   b. $PKR^{*} + ueIF2a_{GDP} + ueIF2a_{GDP} \leftrightarrows PKR^{*} : ueIF2a_{GDP} : ueIF2a_{GDP} \rightarrow PKR^{*} + 2peIF2a_{GDP}$
   $(k_{f} = 1, \; kr = 0.1, \; k_{c} = 1)$
   c. Basal (no stress) phosphorylation: $ueIF2a_{GDP} \rightarrow peIF2a_{GDP} (k_{f} = 0.01)$
   $PKR^{*} + ueIF2a_{GDP} \leftrightarrows PKR^{*} : ueIF2a_{GDP} \rightarrow PKR^{*} + ueIF2a_{GDP}$
   $(k_{f} = 1, \; kr = 0.1, \; k_{c} = 1)$

4. **eIF2B Dynamics:** eIF2B dynamics is included in the models only if the ternary complex is included.
   a. $ueIF2a_{GDP} + eIF2B \leftrightarrows ueIF2a_{GDP} : eIF2B \rightarrow ueIF2a_{GTP} + eIF2B$
   $(k_{f} = 1, \; kr = 0.1, \; k_{c} = 1)$
   $peIF2a_{GDP} + eIF2B \rightarrow peIF2a_{GDP} : eIF2B$
   $(k_{f} = 1)$
   b. $eIF2B_{a} \leftrightarrows eIF2B_{i}$
   $(k_{f} = 0.1, \; kr = 0.1)$
   $ueIF2a_{GDP} + eIF2B_{a} \leftrightarrows ueIF2a_{GDP} : eIF2B_{a} \rightarrow ueIF2a_{GTP} + eIF2B_{a}$
   $(k_{f} = 1, \; kr = 0.1, \; k_{c} = 1)$
   $peIF2a_{GDP} + eIF2B_{a} \rightarrow peIF2a_{GDP} + eIF2B_{i}$
   $(k_{f} = 1)$

5. **ATF4 Translation:**
   a. $\Phi \rightarrow ATF4 \; (k_{f} = 1)$
   b. $\Phi \rightarrow ATF4 \; (k_{f} = \frac{peIF2a}{tot-eIF2a} k)$

c. $\Phi \rightarrow ATF4\ (k_f = \frac{peIF2a}{ueIF2a}k)$

d. $\Phi \rightarrow ATF4\ (k_f = \frac{peIF2a}{K + peIF2a}k)$

e. $\Phi \rightarrow mrnaATF4\ (k_f = 0.01)$

$mrnaATF4 \rightarrow mrnaATF4 + ATF4\ (k_f = 1)$

f. $\Phi \rightarrow mrnaATF4\ (k_f = 0.01)$

$mrnaATF4 \rightarrow mrnaATF4 + ATF4\ (k_f = \frac{peIF2a}{tot-eIF2a}k)$

g. $\Phi \rightarrow mrnaATF4\ (k_f = 0.01)$

$mrnaATF4 \rightarrow mrnaATF4 + ATF4\ (k_f = \frac{peIF2a}{ueIF2a}k)$

h. $\Phi \rightarrow mrnaATF4\ (k_f = 0.01)$

$mrnaATF4 \rightarrow mrnaATF4 + ATF4\ (k_f = \frac{peIF2a}{K + peIF2a}k)$

**6. GADD34 Expression:**

a. $\Phi \rightarrow GADD34\ (k_f = 1)$

b. $\Phi \rightarrow GADD34\ (k_f = \frac{peIF2a}{tot-eIF2a}k)$

c. $\Phi \rightarrow GADD34\ (k_f = \frac{ATF4}{K + ATF4}k)$

d. $\Phi \rightarrow mrnaGADD34\ (k_f = 0.01)$

$mrnaGADD34 \rightarrow mrnaGADD34 + GADD34\ (k_f = 1)$

e. $\Phi \rightarrow mrnaGADD34\ (k_f = \frac{ATF4}{K + ATF4}k)$

$mrnaGADD34 \rightarrow mrnaGADD34 + GADD34\ (k_f = 1)$

f. $\Phi \rightarrow mrnaGADD34\ (k_f = \frac{ATF4}{K + ATF4}k)$

$mrnaGADD34 + ATF4 \leftrightarrows mrnaGADD34{:}ATF4 \rightarrow mrnaGADD34 + GADD34 + ATF4$
$(k_f = 1,\ k_r = 0.1,\ k_c = 1)$

**7. eIF2a Dephosphorylation:**

a. $GADD34 + PP1 \leftrightarrows GADD34{:}PP1\ (k_f = 1,\ k_r = 0.1)$

$GADD34{:}PP1 + peIF2a_{GDP} \leftrightarrows GADD34{:}PP1{:}peIF2a_{GDP} \rightarrow GADD34{:}PP1 + ueIF2a_{GDP}$
$(k_f = 1,\ kr = 0.1,\ k_c = 1)$

If *eIF2B sequestration* is included in the model, then this assumption also includes:

$GADD34{:}PP1 + peIF2a_{GDP}{:}eIF2B \leftrightarrows GADD34{:}PP1{:}peIF2a_{GDP}{:}eIF2B$
$\rightarrow GADD34{:}PP1 + ueIF2a_{GDP} + eIF2B\ (k_f = 1,\ kr = 0.1,\ k_c = 1)$

b. $CReP + PP1 \leftrightarrows CReP{:}PP1\ (k_f = 1,\ k_r = 0.1)$

$CReP:PP1 + peIF2a_{GDP} \leftrightarrows CReP:PP1:peIF2a_{GDP} \rightarrow CReP:PP1 + ueIF2a_{GDP}$
$(k_f = 1,\ kr = 0.1,\ k_c = 1)$
$GADD34 + PP1 \leftrightarrows GADD34:PP1\ (k_f = 1,\ k_r = 0.1)$
$GADD34:PP1 + peIF2a_{GDP} \leftrightarrows GADD34:PP1:peIF2a_{GDP} \rightarrow GADD34:PP1 + ueIF2a_{GDP}$
$(k_f = 1,\ kr = 0.1,\ k_c = 1)$

If *eIF2B sequestration* is included in the model, then this assumption also includes:

$CReP:PP1 + peIF2a_{GDP}:eIF2B \leftrightarrows CReP:PP1:peIF2a_{GDP}:eIF2B$
$\rightarrow CReP:PP1 + ueIF2a_{GDP} + eIF2B\ (k_f = 1,\ kr = 0.1,\ k_c = 1)$
$GADD34:PP1 + peIF2a_{GDP}:eIF2B \leftrightarrows GADD34:PP1:peIF2a_{GDP}:eIF2B$
$\rightarrow GADD34:PP1 + ueIF2a_{GDP} + eIF2B\ (k_f = 1,\ k_r = 0.1,\ k_c = 1)$

### B. Supplementary Tables:

**Table S1:** List of selected 12 models and their underlying hypotheses.

| PKR Activation | ATF4 Translation | TC | eIF2a Phospho | eIF2a Dephospho | eIF2B Dynamics | GADD34 Expression | Number of Species | Number of Parameters | objective (MSE) |
|---|---|---|---|---|---|---|---|---|---|
| d | b | Exc. | c | a | Exc. | c | 17 | 18 | 0.00523 |
| d | b | Exc. | c | b | Exc. | d | 20 | 24 | 0.00403 |
| f | b | Exc. | c | b | Exc. | e | 20 | 25 | 0.00230 |
| g | b | Exc. | c | b | Exc. | c | 19 | 23 | 0.00412 |
| g | c | Exc. | c | a | Exc. | e | 18 | 20 | 0.00409 |
| h | b | Exc. | a | a | Exc. | c | 17 | 18 | 0.00537 |
| h | b | Exc. | c | a | Exc. | e | 18 | 21 | 0.00475 |
| i | c | Exc. | c | a | Exc. | c | 18 | 17 | 0.00531 |
| a | b | Inc. | a | b | b | e | 25 | 32 | 0.00327 |
| c | c | Inc. | c | a | b | c | 22 | 26 | 0.00540 |
| e | c | Inc. | a | b | b | d | 25 | 32 | 0.00386 |
| h | b | Inc. | c | a | a | e | 22 | 28 | 0.00479 |

### C. Supplementary Figures:

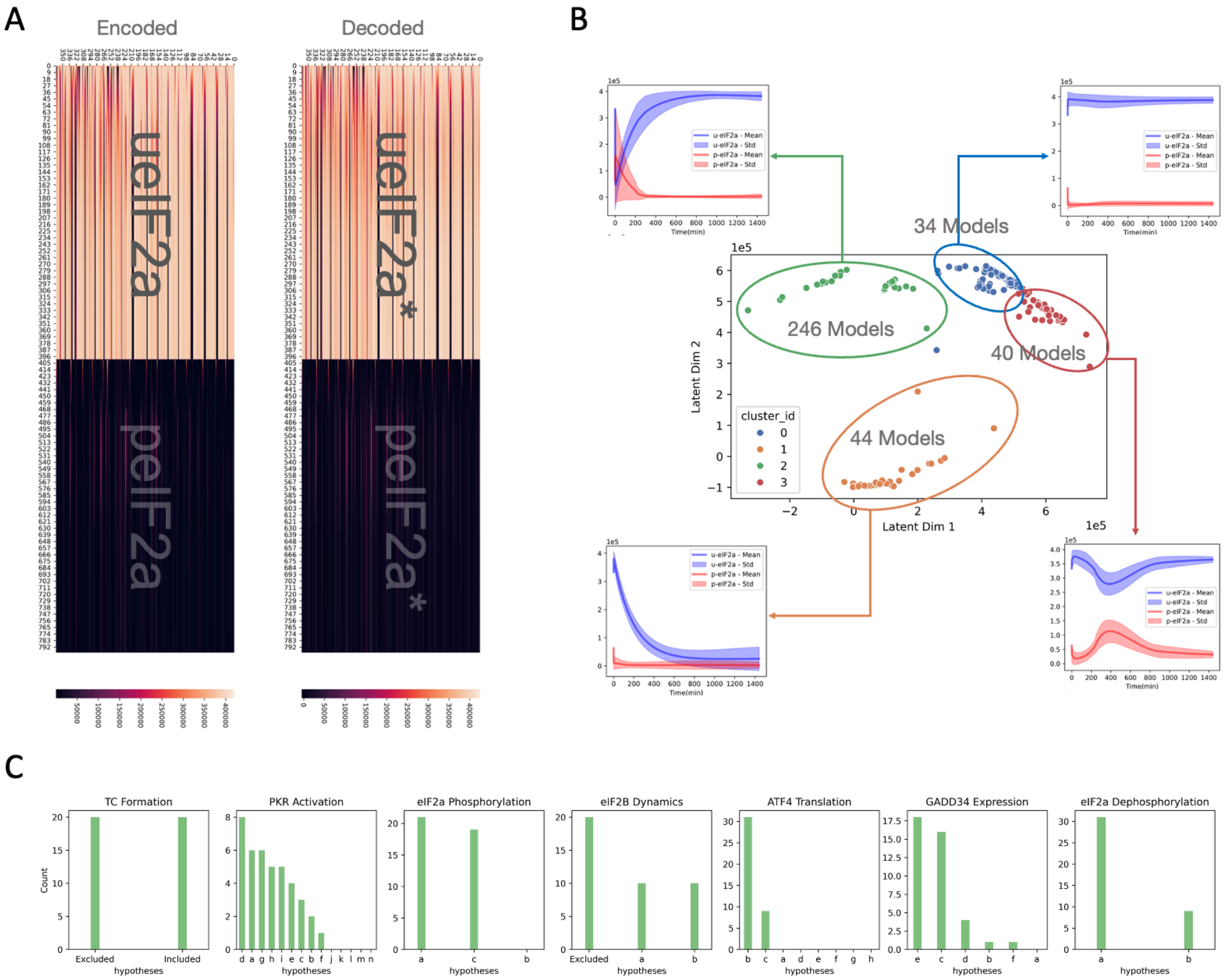


**Figure S1: Model selection via autoencoders: the second iteration with concatenated u-eIF2α, p-eIF2α dynamics.** (A) Encoded and decoded eIF2α dynamics (B) Latent space clusters of models based on eIF2α dynamics (C) Distribution of underlying hypotheses of models in the selected cluster.

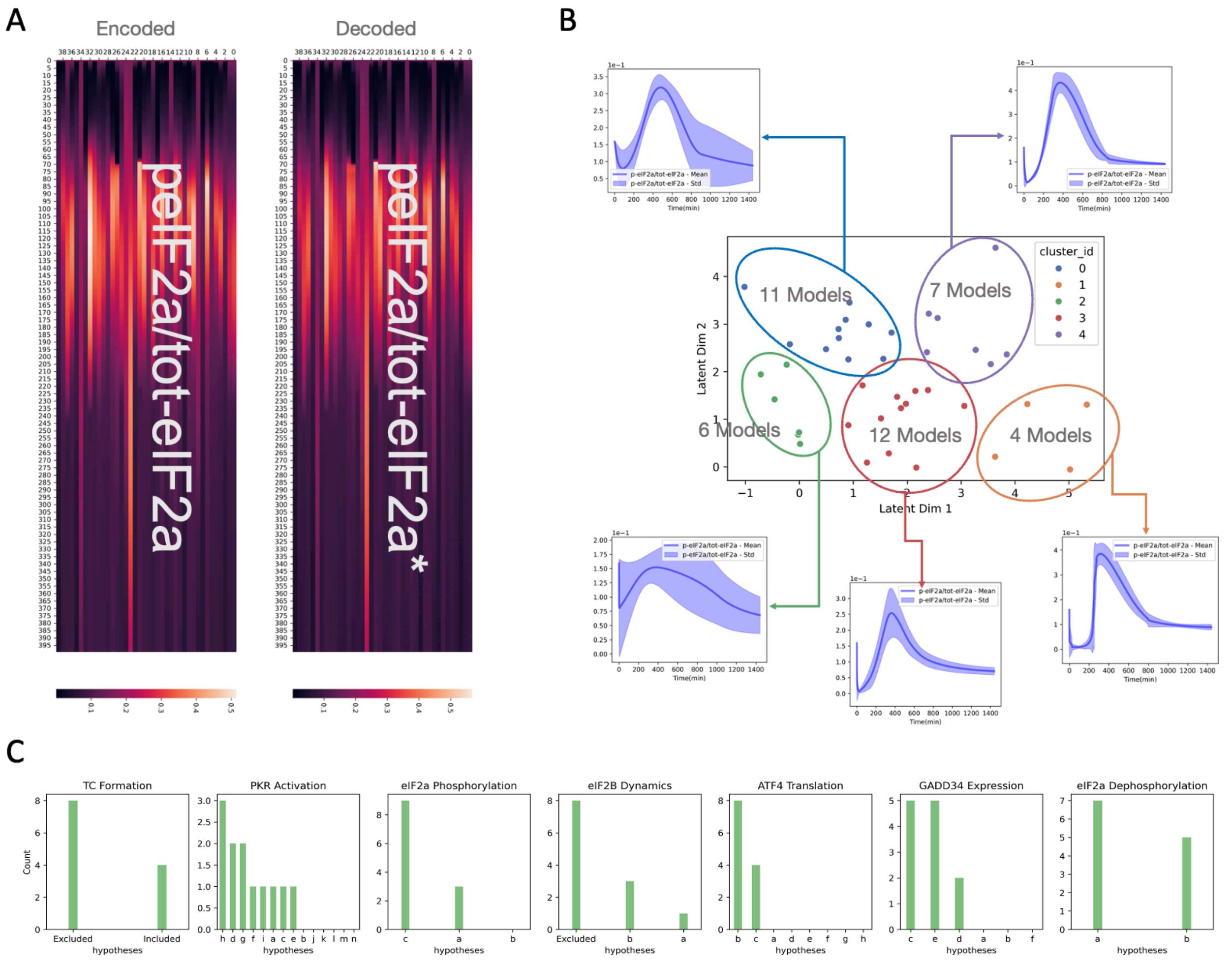


**Figure S2: Model selection via autoencoders: the third iteration with p-eIF2α to total eIF2α ratio over time.** (A) Encoded and decoded p-eIF2α to total eIF2α ratio. (B) Latent space clusters of models based on p-eIF2α to total eIF2α ratio. (C) Distribution of underlying hypotheses of models in the selected cluster.

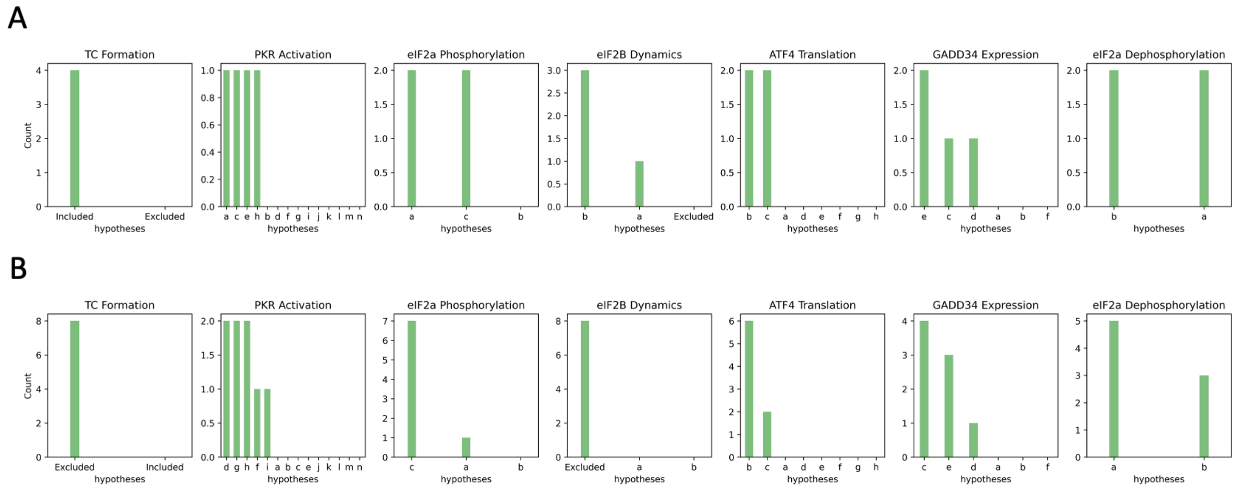


**Figure S3: Distributions of hypotheses in the selected 12 models.** (A) Underlying hypotheses of the large models. (B) Underlying hypotheses of the small models.

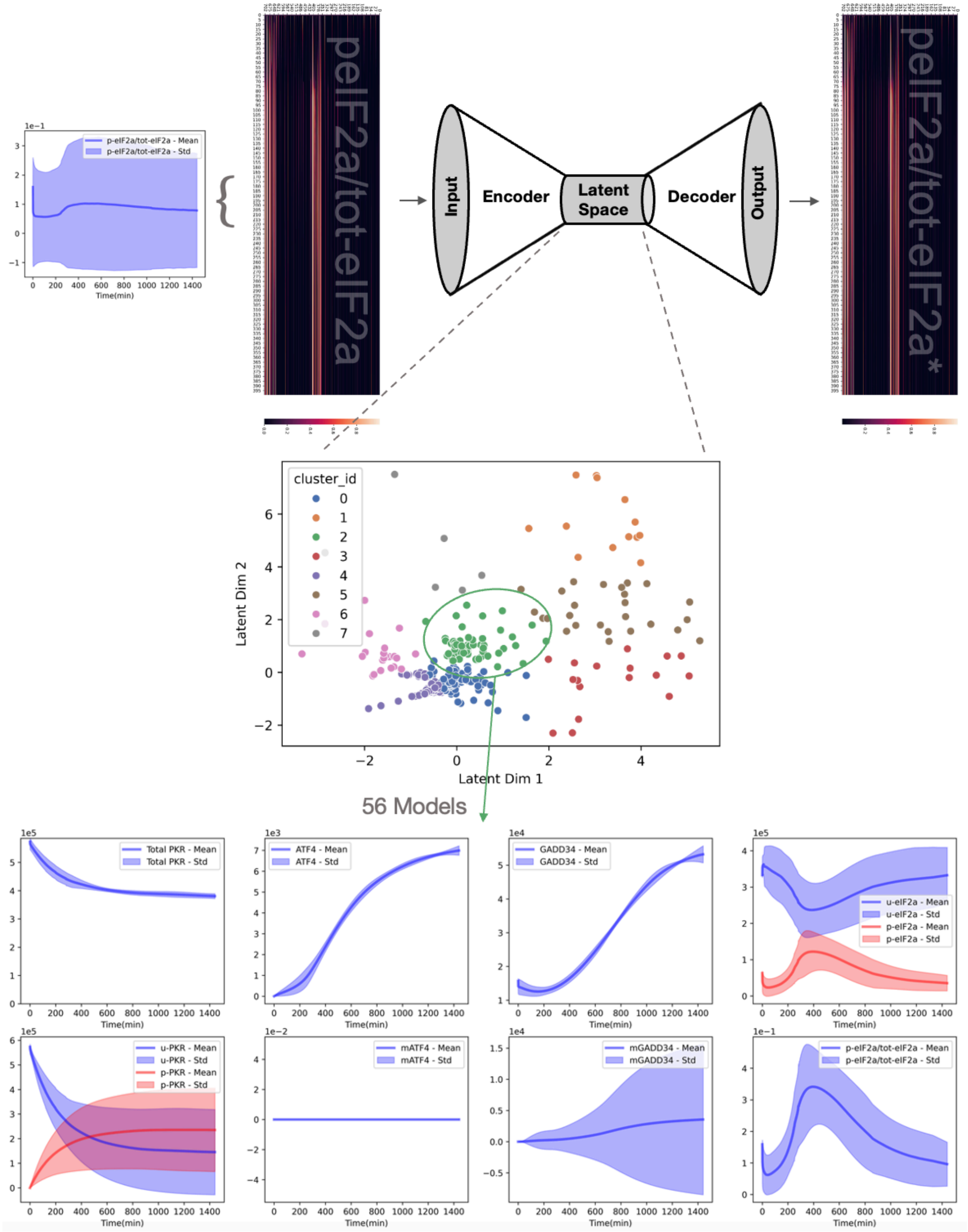


**Figure S4: Robustness analysis of the dynamics embedding approach to the order of iterations.** Here, we start with p-eIF2α to total eIF2α ratio to reduce the number of models. 56 models in the green cluster were selected due to their p-eIF2α to total eIF2α ratio dynamics over time is similar to what was observed before.

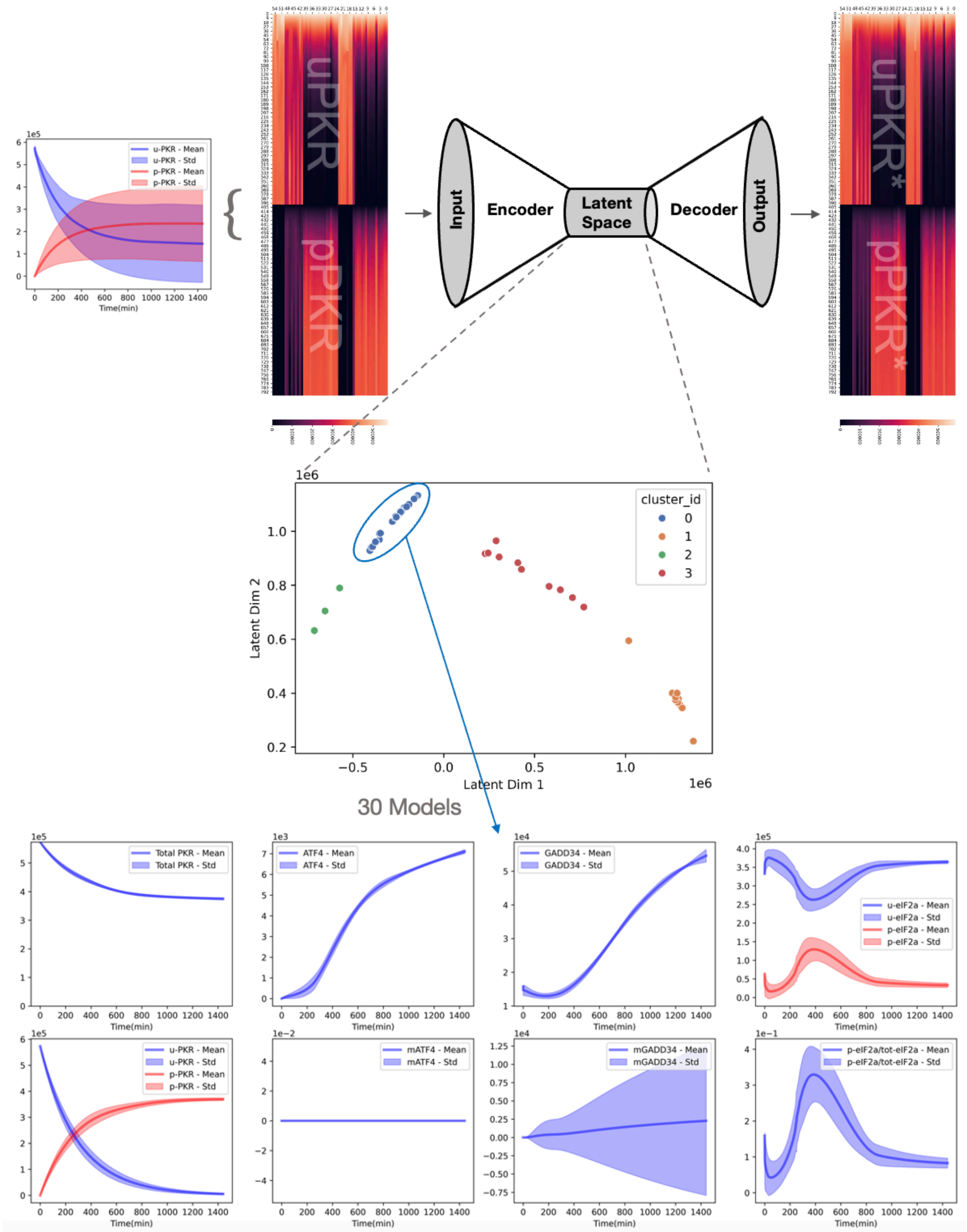


**Figure S5: Robustness analysis of the dynamics embedding approach to the order of iterations.** In the second iteration, we embed the u-PKR and p-PKR dynamics to reduce the number of models. 30 models in the blue cluster were selected based on established physiological constraints.

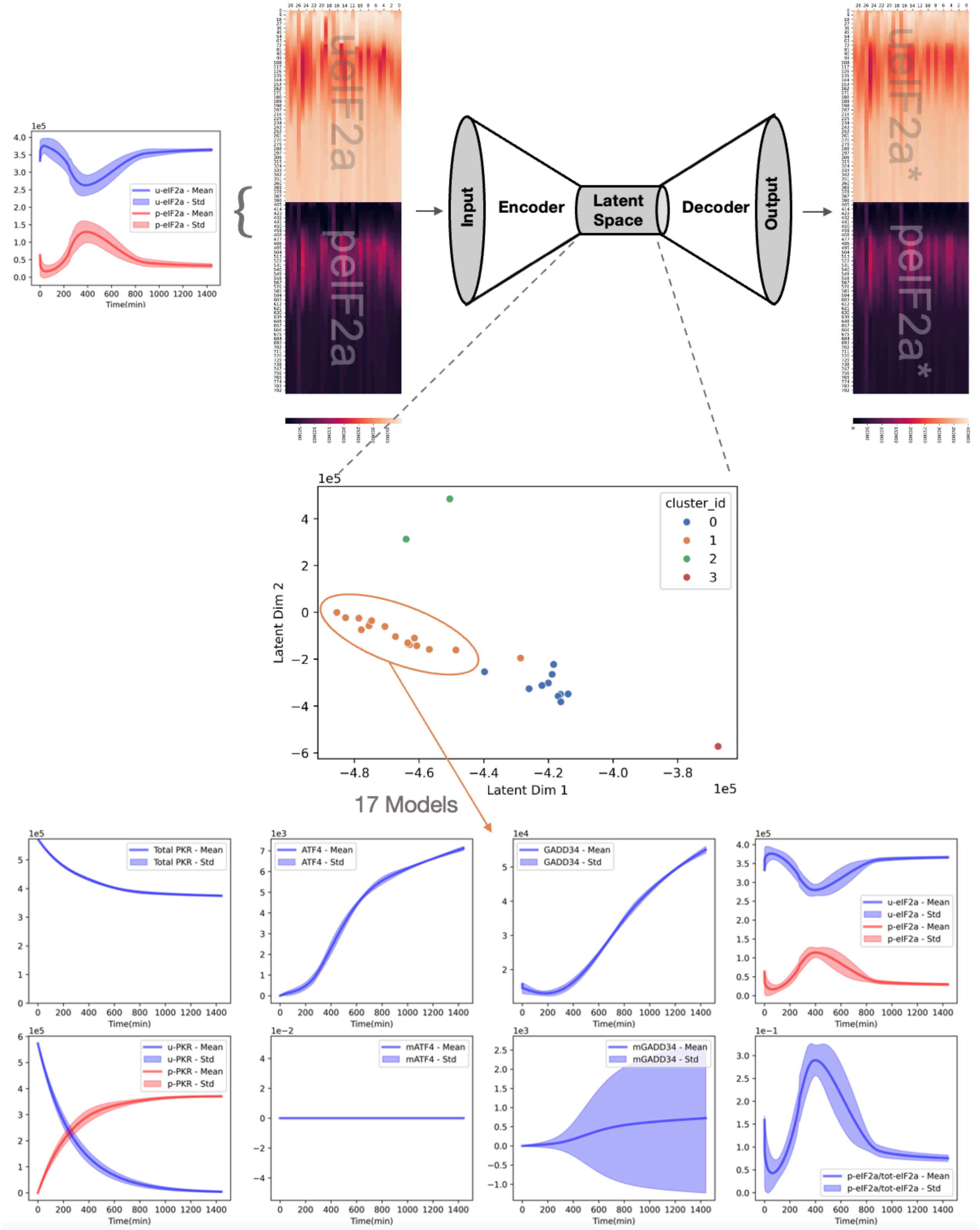


**Figure S6: Robustness analysis of the dynamics embedding approach to the order of iterations.** In the third iteration, we embed the u-eIF2α and p-eIF2α dynamics to reduce the number of models. 17 models in the orange cluster were selected based on established physiological constraints.

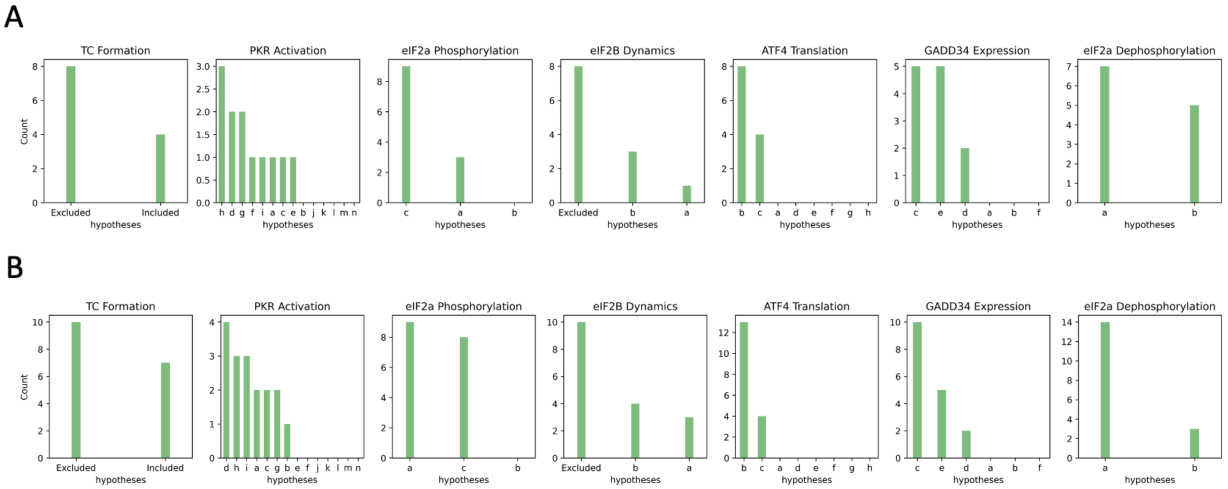


**Figure S7: Comparison of the selected models obtained when different orders of iterations are performed.** (A) Models selected by first embedding PKR dynamics (12 models). (B) Models selected by first embedding p-eIF2α to total eIF2α ratio over time (17 models). Notice that this set of 17 models includes all 12 models identified in our initial analysis.

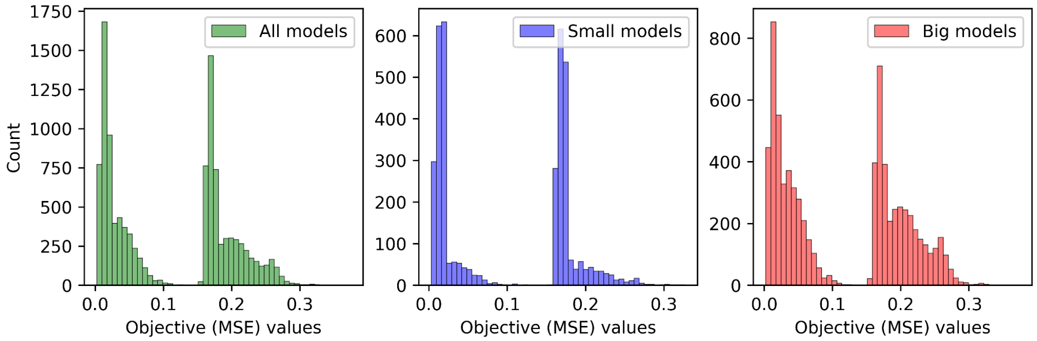


**Figure S8: MSE distributions of all 12,096 models.** The bimodal distribution of MSE values across all mechanistic hypotheses illustrates the fitness landscape: one large peak represents models with poor fitness (high MSE), and a smaller peak/tail represents models that fit the experimental proteomics data well (low MSE).
